# A Model to Simulate the Interference Effects of Acute Aerobic Exercise on Rate of Force Development in Weightlifters

**DOI:** 10.64898/2025.12.18.695222

**Authors:** Emidio E. Pistilli, Alan D. Mizener, Stuart A. Clayton

## Abstract

Compelling data supports the concept that concurrent strength and aerobic training can interfere with adaptations to strength training related to speed, power, and rate of force development (RFD). Studies on this topic have primarily utilized non-athlete participants, with data subsequently extrapolated to athlete populations. Since there may be hesitancy to have athletes involved in research on the interference effects of aerobic exercise on strength specific adaptations, we built a model to simulate these effects and develop an equation to predict interference. The hypothesis tested in this project was that aerobic exercise is the main driver of interference on weightlifting-associated RFD, and that this interference can be modeled to generate an equation to predict changes in RFD. Python software was used to generate the model, perform the simulation and optimize variables that contribute to the interference effect.

A nonlinear exponential interference model was created that simulates the changes in RFD from a force-time curve in response to acute aerobic exercise lasting from 2-minutes to 60-minutes in duration. RFD sensitivity to aerobic exercise was modeled such that values were reduced by approximately 70% after aerobic exercise less than 10 minutes, with further reduction to 80% with durations greater than 10-minutes. The prediction equation can be used by strength coaches to predict the interference of acute aerobic exercise on RFD and subsequently allow the coach to make informed decisions on training program design if there is a need to include aerobic exercise in a strength and/or power-based periodized plan.

## INTRODUCTION

Performing both strength and aerobic exercise training (i.e., concurrent training) is recommended to develop and maintain musculoskeletal and cardiorespiratory fitness, respectively, in healthy adults (1). The expected training-induced adaptations from structured strength training and aerobic exercise training, when performed independently, are well-established and are divergent in nature as described in classic publications (2–4) and in more recent reviews (5–7). The concept of an “interference” on strength training-induced adaptations when performing concurrent strength and aerobic exercise was first suggested by Hickson (8) and has received significant research attention since this initial study. Published research on this topic suggests that concurrent strength and aerobic training may induce minimal interference on training-associated improvements in muscle strength and hypertrophy in untrained people (9–16). However, the potential for interference from aerobic exercise on muscle hypertrophy and strength does become greater in athlete populations, especially for athletes involved in strength sports (17,18). Compelling data also exist suggesting that concurrent strength and aerobic training can interfere with specific adaptations to strength training related to speed, power, and rate of force development (RFD), variables that are associated with optimal nervous system contributions to performance (9–11,19). For example, in their seminal paper, Dudley and Djamil (9) demonstrated that concurrent strength and aerobic training limited the degree of angle-specific maximal torque at fast velocities but not at slow velocities. Hakkinen et al.,(11) also suggest concurrent training interferes with explosive strength performance that is mediated by neural activation of skeletal muscles. When considering the collective adaptations expected from strength training, RFD is likely more sensitive to the inclusion of aerobic exercise than is muscle strength and hypertrophy.

Sports that require speed, power, and RFD include weightlifting and track-and-field events such as sprinting, jumping and throwing. The training programs of these athletes, especially weightlifters, focus almost exclusively on the competition lifts (i.e., snatch; clean and jerk) and weightlifting derivatives to enhance power and RFD (20–23), and rarely include periods of aerobic exercise. To the Authors’ knowledge, no study has been completed in highly trained weightlifters to determine the potential interference effect of acute aerobic exercise on adaptations to weightlifting training and performance. Published studies on this topic utilize healthy subjects that are typically not training for sport performance and conclusions are subsequently extrapolated to athlete populations (9,11–13,17). Given that there may be hesitation to have weightlifters involved in a study to directly determine the interference effects of aerobic exercise on performance, there is a need for an alternative approach to predict or estimate these interference effects. The goal of this project was to create a model to simulate the effects of varying durations of acute aerobic exercise on the subsequent changes to RFD in weightlifters, and to utilize the model to generate an equation that can predict the changes in RFD immediately following acute aerobic exercise. The hypothesis tested in this project was that aerobic exercise is the main driver of interference on weightlifting-associated RFD, and that this interference can be modeled to generate an equation to predict changes in RFD. We built the model such that acute aerobic exercise will cause the slope of a force-time curve from an isometric mid-thigh pull to decrease (i.e., flatten) with increasing durations of acute aerobic exercise. The RFD prediction equation which we have generated allows a weightlifting coach to enter an athlete’s current RFD value, and then estimate the change in RFD, in both absolute loss and percentage loss, following any duration of aerobic exercise. The value in this equation is that it will allow a coach to make informed decisions on acute program design and estimate the potential effects to RFD if aerobic exercise of any duration is performed.

## METHODS

### Experimental Approach to the Problem

The purpose of this project was to develop a predictive equation that can be utilized by coaches and athletes to predict the interference effects of acute aerobic exercise on the force-time curve in weightlifters, specifically predicting changes in RFD that are reflected in the initial slope of the curve. Code to run the simulations was written in Python (version 3.13.7) and is freely available at https://github.com/PistilliLab/RFD_Paper. A model was created to simulate the effects of increasing durations of aerobic exercise on the force-time curve, from a minimum of 2-minutes to a maximum of 60-minutes. The model makes the following assumptions: 1) aerobic exercise duration, defined as “d”, is the main driver of interference on RFD; 2) RFD is sensitive to aerobic exercise such that negative effects can be observed with as little as 2 minutes of aerobic exercise; 3) the rate of RFD decays nonlinearly with increasing durations of aerobic exercise such that greater RFD loss is seen with aerobic exercise lasting <10 minutes and continued smaller reductions in RFD occur with aerobic exercise >10 minutes; 4) while differences in RFD from an isometric mid-thigh pull exist based on sex and weightlifting experience(24–26), maximum RFD in the model is set to 15,000 N to represent the conceptual interference from aerobic exercise.

### Simulation Model Variables

The general form of the prediction equation is as follows:

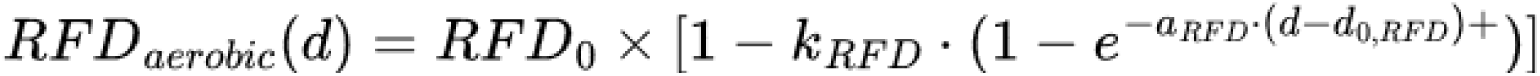

**RFD_₀_** = the athlete’s baseline RFD (i.e., maximum RFD without interference).

**d** = aerobic exercise duration (minutes).

**d**_0,RFD_ = onset time when RFD first begins to be affected.

**αRFD** = decay rate constant (i.e., how quickly interference is observed in response to aerobic exercise.

**κRFD** = RFD Scaling Factor (i.e., sets the maximum that RFD loss occurs with extended duration of aerobic exercise).

### Model Generation

The effect of aerobic exercise duration on RFD was initially modeled with either a linear or nonlinear rate of decay. **Figure 1** displays the general form of absolute and percentage RFD loss when modeled as a linear rate of decay compared to a nonlinear rate of decay. The nonlinear rate of decay model was selected based on the predicted effects of aerobic exercise on the nervous system and sensitivity of RFD to this mode of exercise in weightlifters. As such, greater loss of RFD is seen with aerobic exercise <10-minutes, with continued smaller changes observed with durations of aerobic exercise >10-minutes. The nonlinear rate of decay was subsequently modeled by adjusting “α” to simulate the sensitivity of RFD to the effects of aerobic exercise **(Figure 2)**. Lastly, the nonlinear RFD scaling factor was modeled by adjusting “κ” to establish the limit for RFD loss **(Figure 3)**. Based on these simulations, a nonlinear model was selected with a rate of decay (α) set at a value of 0.3 and an RFD scaling factor (κ) set at 0.8, which would be applicable to weightlifters and other anaerobic athletes training for enhanced RFD.

**Figure 1:**
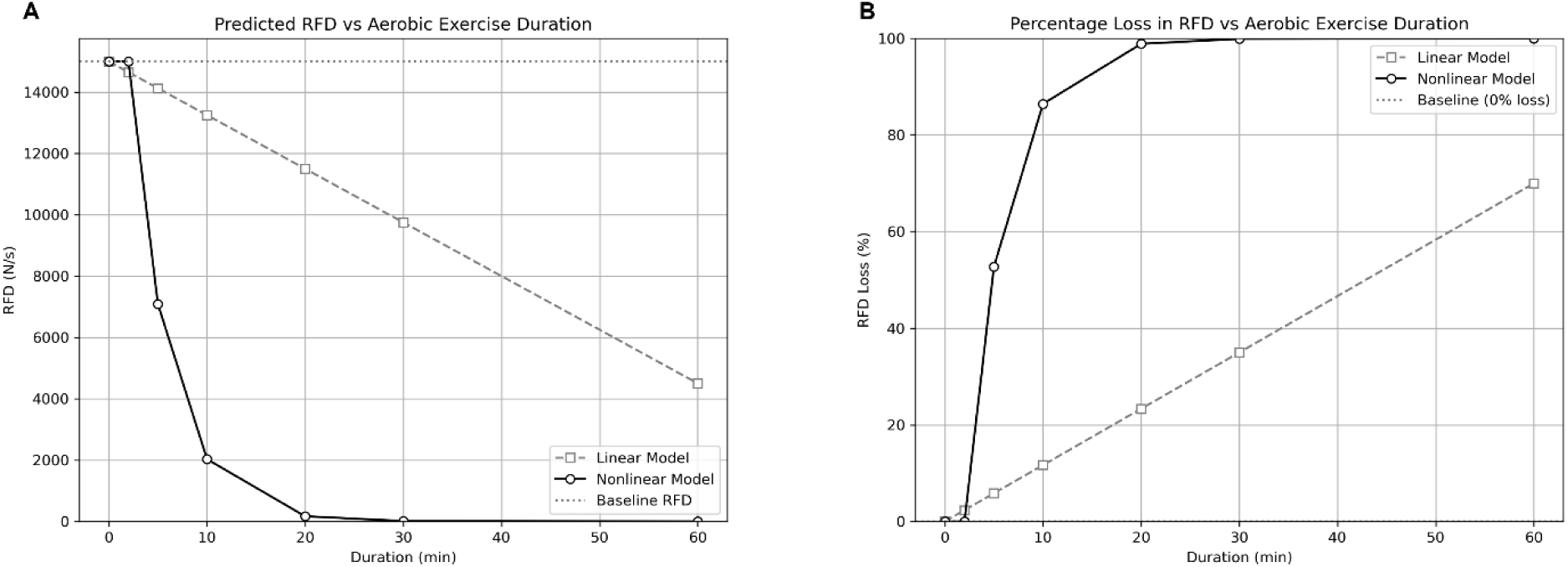
Comparison of linear and nonlinear loss of RFD in response to aerobic exercise. The initial step in developing this model was to determine whether the loss of RFD following acute aerobic exercise would occur as a progressive linear loss or an exponential nonlinear loss. **(A)** The nonlinear loss of absolute RFD produced a curve depicting a rapid decline in response to aerobic exercise <10-minutes in duration, with predicted RFD approaching zero after 30-minutes of aerobic exercise. **(B)** The nonlinear loss of RFD percentage produced a curve that predicted greater than 80% loss in response to 10-minutes of aerobic exercise, with percentage loss approaching 100% loss after 30-minutes of aerobic exercise. While these initial simulations present the unrealistic scenario of an athlete approaching complete loss of RFD, they do support the concept that the exponential nonlinear loss of RFD better models the expected sensitivity of RFD to acute aerobic exercise compared to a progressive linear loss.

**Figure 2:**
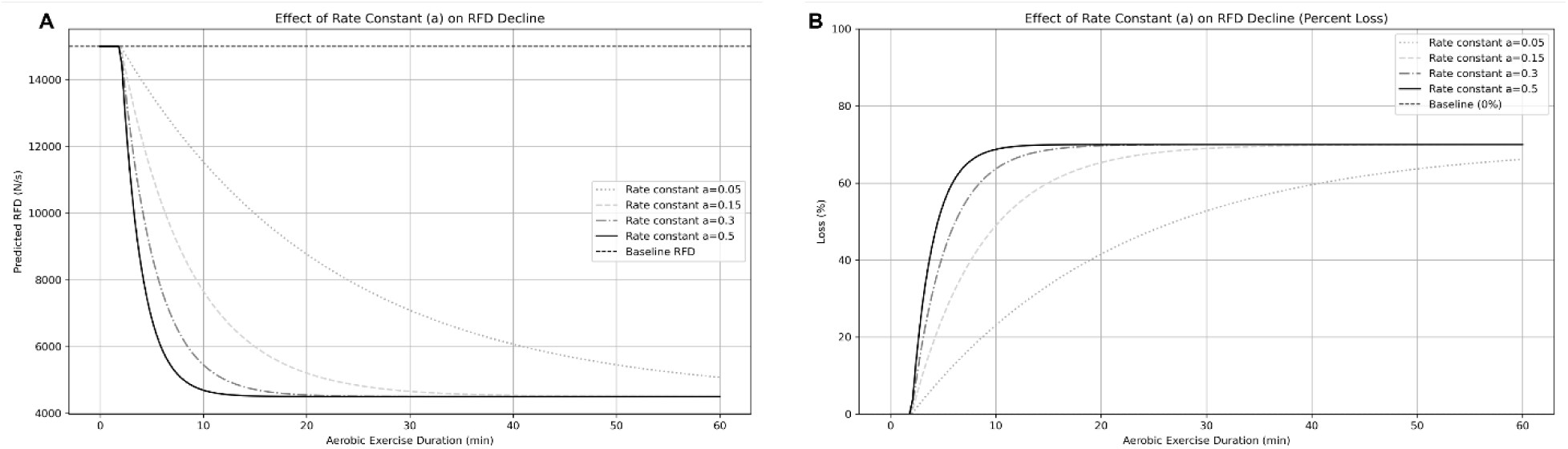
The effects of differing rates of decay on the loss of RFD in response to acute aerobic exercise. RFD is a training-associated variable that is known to be sensitive to the inclusion of aerobic exercise. Therefore, we needed to develop a model to simulate that expected sensitivity. The rate of decay was adjusted to produce a series of curves depicting the absolute loss of RFD in response to acute aerobic exercise. A value of 0.3 was selected to represent the expected sensitivity of RFD to acute aerobic exercise.

**Figure 3:**
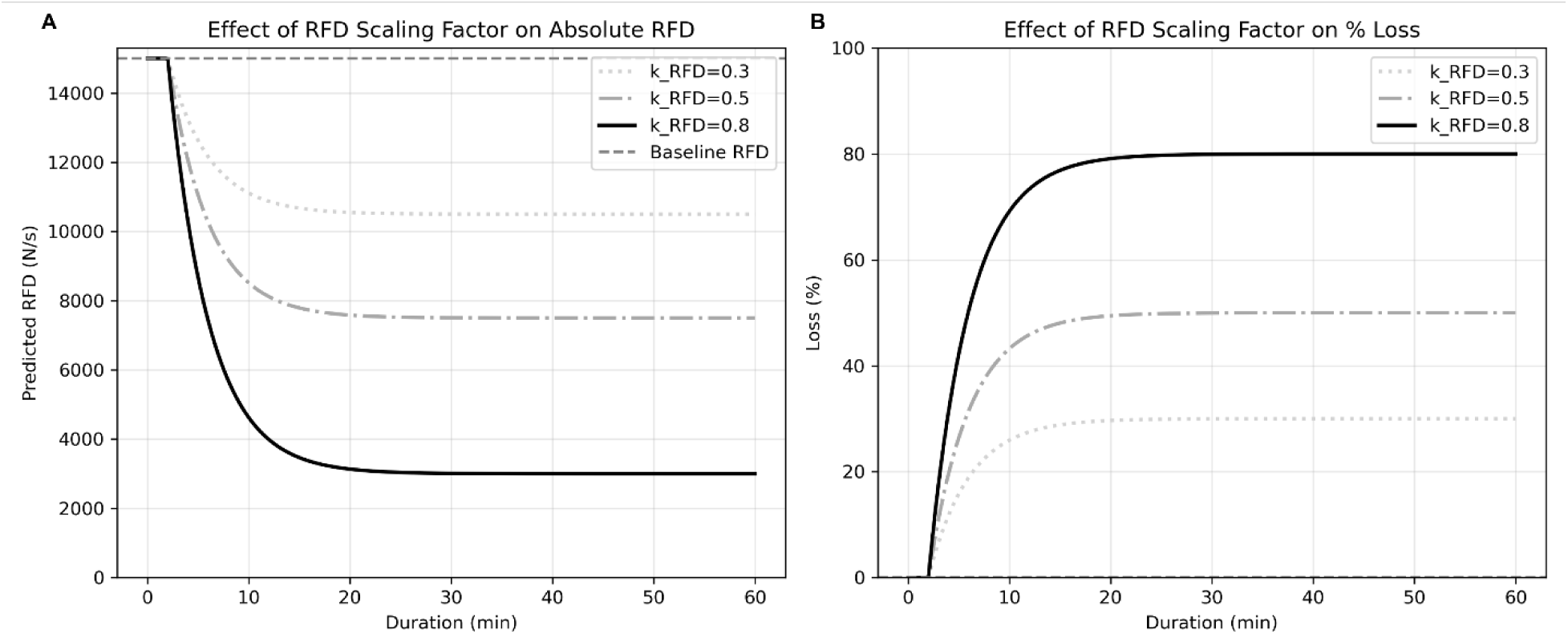
The effects of differing RFD scaling factors on the loss of RFD in response to acute aerobic exercise. While RFD is expected to be sensitive to acute aerobic exercise, it is not expected that an athlete’s RFD would approach zero even after very long durations of aerobic exercise. Therefore, we needed to develop the model to account for an expected maximum loss of RFD. (A) The RFD scaling factor was adjusted to produce a series of curves depicting the absolute loss of RFD following acute aerobic exercise. (B) The RFD scaling factor was adjusted to produce a series of curves depicting the percentage loss of RFD following acute aerobic exercise. A value of 0.8 was selected to represent the theoretical maximum loss of RFD, which would be equivalent to a loss of 80% of an athlete’s maximal RFD.

### Prediction Equation and Code

The RFD prediction equation was further developed into a code that allows coaches to input an athlete’s known peak RFD value and the duration of aerobic exercise to estimate the absolute and percent change in RFD. The resulting RFD value represents the change in slope of the initial phase of the force-time curve when an isometric mid-thigh pull is performed subsequent to the single bout of aerobic exercise. The full interactive code to allow coaches to predict changes in RFD values after aerobic exercise is freely available at https://github.com/PistilliLab/RFDsimulator. In addition, an interactive webapp has been created that allows a coach to enter the above parameters and estimate RFD changes after aerobic exercise (https://rfdsimulator.streamlit.app/). This webapp will allow coaches to enter baseline RFD values, aerobic exercise durations as well as adjust the rate of decay and RFD scaling factor to customize the prediction equation for specific athlete populations.

## RESULTS

### Model Simulation

The nonlinear RFD interference model was tested with aerobic exercise durations in minutes set at: 0; 2; 5; 10; 20; 30; and 60. The simulation produced a series of force-time curves that represent an isometric mid-thigh pull after completing the different durations of aerobic exercise **(Figure 4)**. Absolute loss and the percentage loss of RFD was subsequently predicted from these force-time curves based on an initial maximum value for RFD of 15,000 N s^-1^, a rate of decay of 0.30, and a maximal RFD scaling factor of 0.8. **Figure 5A** shows the estimated changes in absolute RFD with increasing durations of aerobic exercise, such that RFD begins to drop following 2-minutes of aerobic exercise. Furthermore, the steep portion of the absolute RFD loss curve occurs with <10-minutes of aerobic exercise, with smaller and less dramatic changes with >10-minutes of aerobic exercise, reflecting the sensitivity of RFD to aerobic exercise. **Figure 5B** shows the estimated percentage loss of RFD with increasing durations of aerobic exercise. The steep portion of percentage loss of RFD curve occurs with <10-minutes of aerobic exercise, such that approximately 60-70% of baseline RFD is lost. With aerobic exercise >10-minutes, the percentage loss of RFD continues to a maximum of ∼80% of baseline RFD, based on the RFD scaling factor.

**Figure 4:**
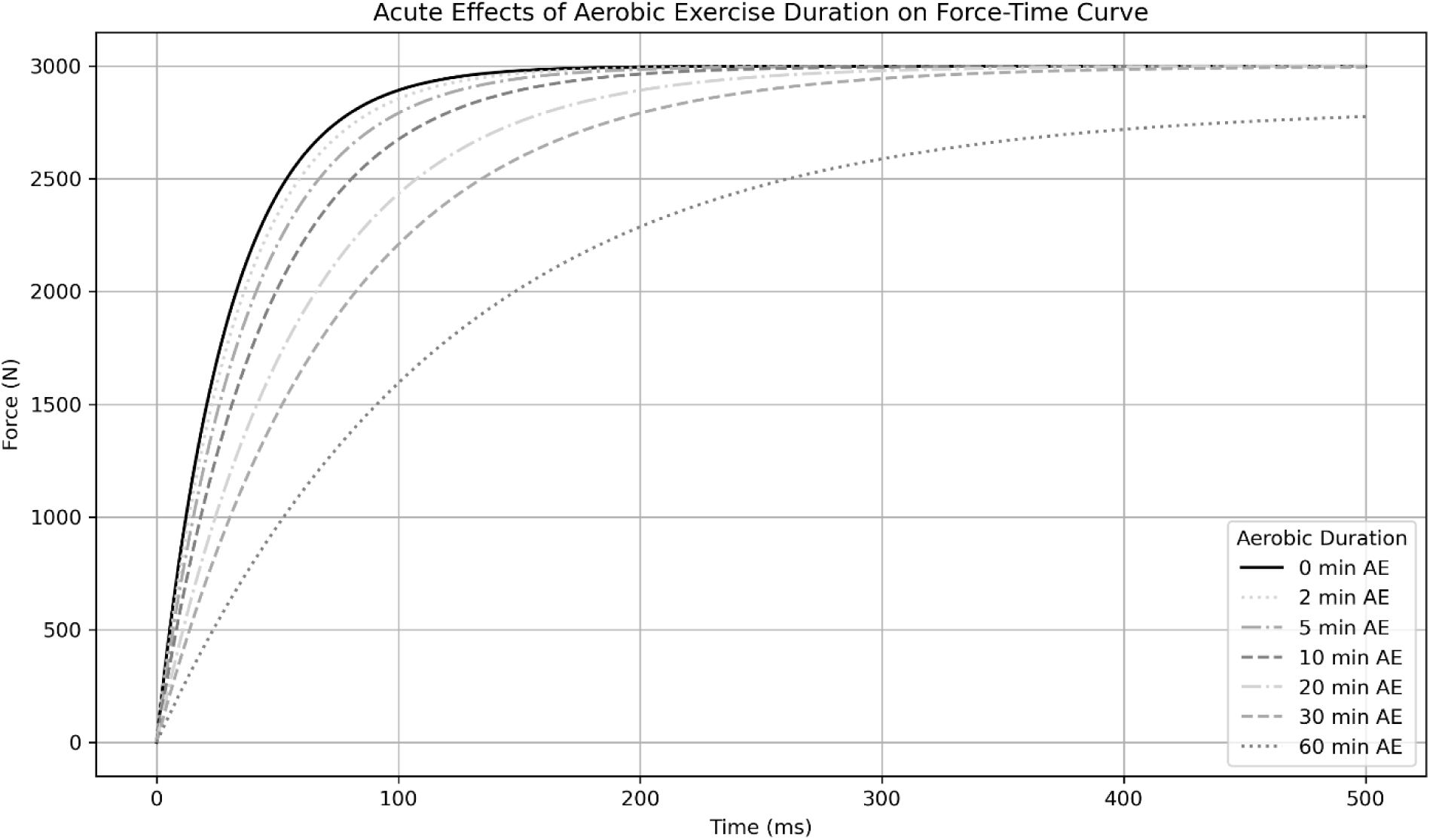
The effects of acute aerobic exercise on the force-time curve. The simulation model was used to estimate the changes in the force-time curve produced from an isometric mid-thigh pull following various durations of aerobic exercise. A series of force-time curves were generated in response to increasing duration of aerobic exercise, from 5-minutes to 60-minutes, and can be compared to the force-time curve following no aerobic exercise (blue line). The changes in the force-time curves were produced using a rate of decay value of 0.3 and an RFD scaling factor of 0.8.

**Figure 5:**
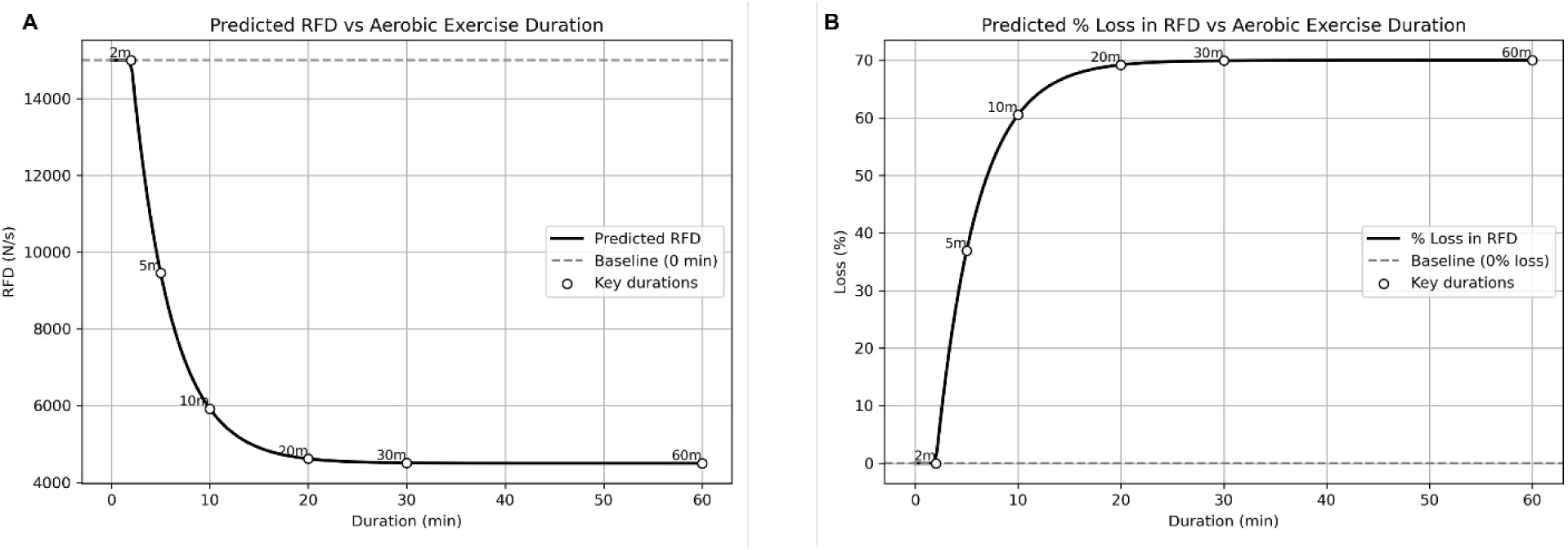
The final model depicting the estimated loss of RFD in response to various durations of acute aerobic exercise. The final model parameters used to generate these curves were a rate of decay value of 0.3, an RFD scaling factor of 0.8, and the assumption that RFD loss would be minimal following aerobic exercise <2-minutes. **(A)** The absolute loss of RFD in response to acute aerobic exercise. The rapid initial loss of RFD occurs in response to aerobic exercise durations <10-minutes, with smaller losses that may plateau following aerobic exercise >30-minutes. **(B)** The percentage loss of RFD in response to acute aerobic exercise. The rapid initial loss of RFD approaches 70% in response to aerobic exercise durations <10-minutes, with smaller losses that approaching 80% following aerobic exercise >30-minutes. **(C)** Estimates of absolute loss and percentage loss of RFD in response to varying durations of aerobic exercise.

### Coach Tunable Variables

The Python code allows a coach to adjust the variables based on the specific athletes that may be tested. After entering an athlete’s absolute RFD value, the following variables can be adjusted: aerobic exercise duration (d); rate of decay (a); and RFD scaling factor (k). The model produces an RFD loss curve based on duration of aerobic exercise from 2-minutes to 60-minutes. A coach would be able to enter in any aerobic exercise duration within this range and receive an estimate of the interference on RFD, reflecting any situation in which an athlete may need to perform aerobic exercise (i.e., rehabilitation; conditioning; etc.). The rate of decay is an estimate of the sensitivity of RFD to aerobic exercise. Therefore, the coach can adjust the rate of decay to simulate changes to RFD in athletes that may be more or less susceptible to the interference of aerobic exercise **(Figure 2)**. Lastly, the RFD scaling factor can be adjusted to set the maximum loss of RFD due to aerobic exercise interference to estimate these changes in numerous types of athletes **(Figure 3)**.

## DISCUSSION

The basic principles of specificity and overload provide the framework for training program design. Strength & Conditioning Coaches utilize these two principles to manipulate training volume and intensity throughout the course of a program in order to peak athletes for competitions. This is the basis for the concept of Periodization, defined as a “theoretical and practical construct that allows for the systematic, sequential, and integrative programming of training interventions into mutually dependent periods of time in order to induce specific physiological adaptations that underpin performance outcomes” (27). Training volume and intensity are manipulated in the training of athletes to enhance specific performance variables such as maximum force output, RFD, and power. RFD is specific to athletes that require maximal force output in very short periods of time, such as weightlifters (23,26,28–30), is based on the ability of the nervous system to rapidly activate skeletal muscles and can be enhanced with specific training methods (11,19,20,31–33). A significant body of literature exists to support the concept that aerobic exercise can have a negative effect on aspects of nervous system function, which is tightly connected to RFD, power, and speed (9–11,19). Therefore, athletes and coaches should be aware of this potential interference effect when designing training programs for athletes that require peak RFD and consider the risk-to-benefit ratio of including aerobic exercise in the training program for these athletes.

It is our assumption that coaches involved in the training of weightlifters would be hesitant to allow their athletes to be involved in a research study to directly test the interference effect of aerobic exercise on performance, which was the reason to conduct this simulation. To highlight how this prediction equation could be utilized, we can refer to a prior publication that described performance variables in weightlifters across 20 weeks of training (25). This study utilized active and competitive male and female national caliber weightlifters. Isometric mid-thigh pull tests were performed, as part of a laboratory test battery, during training weeks 1, 6, 10, 13, 17, and 20, associated with different phases of periodized training. At week 1, the male weightlifters had a group average RFD value of 16,652 N s^-1^ and the female weightlifters had a group average RFD value of 7663 N s^-1^. We can estimate the absolute RFD and percentage RFD loss in these groups using the new RFD interference model with the following variables: rate of decay set at 0.3; RFD scaling factor set at a maximum of 0.8; duration of aerobic exercise set at 10 minutes. In this example, the estimated RFD would be reduced by 72.7% following 10-minutes of aerobic exercise, which would be 4538.9 N s^-1^ for the male weightlifters and 2088.7 N s^-1^ for the female weightlifters.

The idea that training should be based on the movement patterns of athletes and that muscles involved in those movements should be targeted for strengthening has been evident for at least 70 years. Jim Murray and Peter Karpovich stated in 1956 that “any competent coach should be able to make valuable use of weight training by studying the action of his players’ bodies in specific sports, and then selecting resistance exercises which strengthen the muscle groups involved” (34). As stated by Gary Dudley in 1985, “obviously, the nature of the adaptative response to training is specific to the training stimulus” (9,10). In the study by Dudley and Djamil (9), recreationally trained people were randomly assigned to 1 of 3 groups that performed exercise training for 7 weeks: an aerobic training only group; a strength training only group; a concurrent training group that performed both of the aerobic and strength training programs. Strength training was performed on an isokinetic device in which two 30s sets of leg extensions were performed at an angular velocity of 4.19 rad s^-1^. At the end of the 7 weeks, there were no differences in improvement of aerobic power when comparing the aerobic training group and the concurrent training group, suggesting that concurrent training does not interfere with improvements in aerobic capacity and subsequent studies have confirmed this observation (11–13,15,17). To assess isokinetic strength, participants were tested for maximal torque at a range of velocities up to 5.03 rad sec^-1^. Participants in the strength training group improved torque at all velocities tested that were equal to and below the velocity in which training occurred (i.e., 4.19 rad sec^-1^). However, there were no improvements in torque above 4.19 rad sec^-1^. Interestingly, participants in the concurrent training group only improved torque at the three lowest velocities tested, while no improvements were observed at the velocity in which training occurred. There are two critical conclusions to be taken from the results of this study. First, the specificity of training can be observed in the strength training group that completed the 7 weeks of strength training at 4.19 rad.sec^-1^. Therefore, athletes that must produce muscle contractions at high velocities need to include training modalities at or near the velocity required for sport performance. Second, the inclusion of the aerobic exercise training utilized in this study interfered with training-associated improvements at fast velocities of training, and improvements were only observed at the lowest velocities tested. Extending these results to sport performance, it may be concluded that aerobic exercise in the training program of an anaerobic athlete has the potential to limit force production at fast velocities. This has obvious implications for specific athlete populations that require differing expressions of muscle strength to be successful and any athlete that needs to produce rapid muscle contractions against a resistance should limit exposure to aerobic exercise, as there is the potential to limit RFD and power adaptations to training.

Enhancing RFD requires specific training methods to affect the nervous system. One specific adaptation that occurs is by-passing the orderly recruitment of motor units, a process known as the Henneman Size Principle (35). Motor units in skeletal muscle range in size and the recruitment threshold needed to stimulate the motor unit and associated muscle fibers. In general, small motor units are associated with a greater percentage of type I fibers, have small recruitment thresholds which allow them to be easily recruited, and are associated with lower force output. In contrast, large motor units are associated with a greater percentage of type II fibers, have larger recruitment thresholds to overcome for stimulation, and are associated with large force outputs. The nervous system is organized to recruit the low threshold motor units first and then progressively recruit larger motor units as greater force is required (36,37). To facilitate improvements in RFD, the nervous system can by-pass this orderly recruitment pattern, engage the large motor units initially and take advantage of the greater force potential early in the movement pattern (31,38). As seen in the work from Del Vecchio (19), training for maximal force production and fast contraction speeds are mutually exclusive and increases in RFD are due more to the ability to increase the recruitment speed of motor units than to the amount of muscle mass that is producing that force. Given the numerous reports that demonstrate that aerobic exercise interferes with speed, power, and nervous system adaptations (9,11–13,17), athletes that require high RFD such as weightlifters should perform as little aerobic exercise as possible. Reconsidering the classic literature on mechanisms of aerobic exercise adaptations and how those may impact anaerobic training adaptations is important when strength coaches develop overall training programs for their athletes.

There are a few limitations to this study that should be acknowledged. First, the model was built on the assumptions identified in the methods section and therefore may not address situations outside of these a priori assumptions. For example, the rate of decay used in the model was selected based on our interpretation of the literature on the effects of aerobic exercise on nervous system function and the sensitivity of RFD to this mode of exercise.

However, the rate of decay can be adjusted in the model based on each coach’s interpretation of this effect. Second, we hypothesized that aerobic exercise duration was the main driver of the interference on RFD. But we cannot determine how residual fatigue from acute aerobic exercise on other body systems may impact these RFD estimates. Additionally, we did not model the potential interference of specific types of aerobic exercise, such as running compared to cycling. Finally, the interference estimated in this model reflects how the RFD may be affected when the isometric mid-thigh pull is performed relatively soon after acute aerobic exercise, within 1-2 hours. The model is not able to provide information on how soon RFD would be recovered after completing the acute aerobic exercise or how RFD would be affected with periods of chronic aerobic exercise.

In conclusion, we were able to produce an equation that can estimate the interference of various durations of acute aerobic exercise on subsequent RFD from an isometric mid-thigh pull. This prediction equation and model is specific to weightlifters, but the variables included in the equation can be adjusted for other populations of athletes. The simulation model suggests that RFD may be negatively impacted after as little as 2-minutes of aerobic exercise and that 60%-70% of the maximal RFD may be lost after 10-minutes of aerobic exercise. However, any duration of aerobic exercise, up to 60-minutes, can be tested with this model. The prediction equation can be used by strength coaches to predict the interference of acute aerobic exercise on RFD and subsequently allow the coach to make informed decisions on training program design if there is a need to include aerobic exercise in a strength and/or power-based periodized plan.

## PRACTICAL APPLICATIONS

Coaches that work with weightlifters should consider the tissue and system-level adaptations associated with chronic aerobic and strength training and the potential for aerobic exercise to interference with certain performance variables. Generally, it is recommended to limit aerobic exercise in the overall training program of weightlifters and possibly other anaerobic athletes that train for RFD. While every athlete is different and the performance needs of sports vary, the general conclusion of published studies suggests that aerobic exercise may induce sub-optimal or maladaptive responses to training which are necessary to be successful in the athletes’ respective sports. This view of sport training is very different from the perspective of exercising for health and wellness, with recommendations to perform both aerobic and resistance exercise sessions as part of a total exercise program. In this case, the goal of improving health and wellness supersedes the potential interference of performing aerobic and strength training together. When training athletes for improvements in sports performance, the specific variables that contribute to success in sport must be the priority and a coach must keep the training priorities in mind when designing and implementing training programs. If an athlete requires maximum speed, power output, or RFD, then the training program must focus on enhancing those variables and any mode of exercise that is counter to improving those variables should be excluded from the program. If a situation arises that may require short-term periods of aerobic exercise, this simulation model could be used to estimate the degree of interference of acute aerobic exercise and, subsequently allow the coach to make informed decisions on training program design.

## ACKNOWLEDGEMENTS

Emidio Pistilli is funded through the NIH and NIAMS (R01AR0779445). The Authors report no conflicts of interest. The results of the present study do not constitute endorsement of any product by the authors or the NSCA.

## REFERENCES

1. Garber, C. E., Blissmer, B., Deschenes, M. R., Franklin, B. A., Lamonte, M. J., Lee, I. M., Nieman, D. C., Swain, D. P., and American College of Sports, M. (2011) American College of Sports Medicine position stand. Quantity and quality of exercise for developing and maintaining cardiorespiratory, musculoskeletal, and neuromotor fitness in apparently healthy adults: guidance for prescribing exercise. Med Sci Sports Exerc 43, 1334–1359

2. Saltin, B. (1969) Physiological effects of physical training. Med. Sci. Sports 1, 50–56

3. Pollock, M. L. (1973) The quantification of endurance training programs. Exerc Sport Sci Rev 1, 155–188

4. Atha, J. (1981) Strengthening muscle. Exerc Sport Sci Rev 9, 1–73

5. Grgic, J., Schoenfeld, B. J., Davies, T. B., Lazinica, B., Krieger, J. W., and Pedisic, Z. (2018) Effect of Resistance Training Frequency on Gains in Muscular Strength: A Systematic Review and Meta-Analysis. Sports Med 48, 1207–1220

6. Zouita, A., Darragi, M., Bousselmi, M., Sghaeir, Z., Clark, C. C. T., Hackney, A. C., Granacher, U., and Zouhal, H. (2023) The Effects of Resistance Training on Muscular Fitness, Muscle Morphology, and Body Composition in Elite Female Athletes: A Systematic Review. Sports Med 53, 1709–1735

7. Jones, A. M., and Carter, H. (2012) The effect of endurance training on parameters of aerobic fitness. Sports Med 29, 373–386

8. Hickson, R. C. (1980) Interference of strength development by simultaneously training for strength and endurance. Eur J Appl Physiol Occup Physiol 45, 255–263

9. Dudley, G. A., and Djamil, R. (1985) Incompatibility of endurance-and strength-training modes of exercise. J Appl Physiol (1985) 59, 1446–1451

10. Dudley, G. A., and Fleck, S. J. (1987) Strength and endurance training. Are they mutually exclusive? Sports Med 4, 79–85

11. Hakkinen, K., Alen, M., Kraemer, W. J., Gorostiaga, E., Izquierdo, M., Rusko, H., Mikkola, J., Hakkinen, A., Valkeinen, H., Kaarakainen, E., Romu, S., Erola, V., Ahtiainen, J., and Paavolainen, L. (2003) Neuromuscular adaptations during concurrent strength and endurance training versus strength training. Eur J Appl Physiol 89, 42–52

12. Glowacki, S. P., Martin, S. E., Maurer, A., Baek, W., Green, J. S., and Crouse, S. F. (2004) Effects of resistance, endurance, and concurrent exercise on training outcomes in men. Med Sci Sports Exerc 36, 2119–2127

13. Lee, M. J., Ballantyne, J. K., Chagolla, J., Hopkins, W. G., Fyfe, J. J., Phillips, S. M., Bishop, D. J., and Bartlett, J. D. (2020) Order of same-day concurrent training influences some indices of power development, but not strength, lean mass, or aerobic fitness in healthy, moderately-active men after 9 weeks of training. PLoS One 15, e0233134

14. Wilson, J. M., Marin, P. J., Rhea, M. R., Wilson, S. M., Loenneke, J. P., and Anderson, J. C. (2012) Concurrent training: a meta-analysis examining interference of aerobic and resistance exercises. J Strength Cond Res 26, 2293–2307

15. Sedano, S., Marin, P. J., Cuadrado, G., and Redondo, J. C. (2013) Concurrent training in elite male runners: the influence of strength versus muscular endurance training on performance outcomes. J Strength Cond Res 27, 2433–2443

16. Callister, R., Shealy, M. J., Fleck, S. J., and Dudley, G. A. (1988) Performance adaptations to sprint, endurance and both modes of training. J Appl Sport Sci Res 2, 46–51

17. Kraemer, W. J., Patton, J. F., Gordon, S. E., Harman, E. A., Deschenes, M. R., Reynolds, K., Newton, R. U., Triplett, N. T., and Dziados, J. E. (1995) Compatibility of high-intensity strength and endurance training on hormonal and skeletal muscle adaptations. J Appl Physiol (1985) 78, 976-989

18. Schoenfeld, B. J. (2021) Science and Development of Muscle Hypertrophy, 2 ed., Human Kinetics, Champaign, IL

19. Del Vecchio, A., Casolo, A., Dideriksen, J. L., Aagaard, P., Felici, F., Falla, D., and Farina, D. (2022) Lack of increased rate of force development after strength training is explained by specific neural, not muscular, motor unit adaptations. J Appl Physiol (1985) 132, 84-94

20. James, L. P., Suchomel, T. J., Comfort, P., Haff, G. G., and Connick, M. J. (2022) Rate of Force Development Adaptations After Weightlifting-Style Training: The Influence of Power Clean Ability. J Strength Cond Res 36, 1560–1567

21. Suchomel, T. J., McKeever, S. M., and Comfort, P. (2020) Training With Weightlifting Derivatives: The Effects of Force and Velocity Overload Stimuli. J Strength Cond Res 34, 1808–1818

22. Suchomel, T. J., and Sole, C. J. (2017) Force-Time-Curve Comparison Between Weight-Lifting Derivatives. Int J Sports Physiol Perform 12, 431–439

23. Takei, S., Kambayashi, S., Katsuge, M., Okada, J., and Hirayama, K. (2024) Portions of the force-velocity relationship targeted by weightlifting exercises. Sci Rep 14, 31021

24. Haff, G. G., Carlock, J. M., Hartman, M. J., Kilgore, J. L., Kawamori, N., Jackson, J. R., Morris, R. T., Sands, W. A., and Stone, M. H. (2005) Force-time curve characteristics of dynamic and isometric muscle actions of elite women olympic weightlifters. J Strength Cond Res 19, 741–748

25. Hornsby, W. G., Gentles, J. A., MacDonald, C. J., Mizuguchi, S., Ramsey, M. W., and Stone, M. H. (2017) Maximum Strength, Rate of Force Development, Jump Height, and Peak Power Alterations in Weightlifters across Five Months of Training. Sports (Basel) 5

26. Suarez, D. G., Mizuguchi, S., Hornsby, W. G., Cunanan, A. J., Marsh, D. J., and Stone, M. H. (2019) Phase-Specific Changes in Rate of Force Development and Muscle Morphology Throughout a Block Periodized Training Cycle in Weightlifters. Sports (Basel) 7

27. Haff, G. G., and Triplett, N. T. (eds). (2016) Essentials of Strength Training and Conditioning, Human Kinetics, Illinois

28. Garhammer, J. (1980) Power production by Olympic weightlifters. Med Sci Sports Exerc 12, 54–60

29. Stone, M. H., Sands, W. A., Pierce, K. C., Carlock, J., Cardinale, M., and Newton, R. U. (2005) Relationship of maximum strength to weightlifting performance. Med Sci Sports Exerc 37, 1037–1043

30. Storey, A., and Smith, H. K. (2012) Unique aspects of competitive weightlifting: performance, training and physiology. Sports Med 42, 769–790

31. Nardone, A., Romano, C., and Schieppati, M. (1989) Selective recruitment of high-threshold human motor units during voluntary isotonic lengthening of active muscles. J Physiol 409, 451–471

32. Cormie, P., McGuigan, M. R., and Newton, R. U. (2010) Influence of strength on magnitude and mechanisms of adaptation to power training. Med Sci Sports Exerc 42, 1566–1581

33. Cormie, P., McGuigan, M. R., and Newton, R. U. (2011) Developing maximal neuromuscular power: Part 1--biological basis of maximal power production. Sports Med 41, 17–38

34. Murray, J., and Karpovich, P. V. (1956) Weight Training In Athletics, Prentice-Hall, Inc., Englewood Cliffs, N.J.

35. Henneman, E. (1957) Relation between size of neurons and their susceptibility to discharge. Science 126, 1345–1347

36. McPhedran, A. M., Wuerker, R. B., and Henneman, E. (1965) Properties of Motor Units in a Heterogeneous Pale Muscle (M. Gastrocnemius) of the Cat. J Neurophysiol 28, 85–99

37. McPhedran, A. M., Wuerker, R. B., and Henneman, E. (1965) Properties of Motor Units in a Homogeneous Red Muscle (Soleus) of the Cat. J Neurophysiol 28, 71–84

38. ter Haar Romeny, B. M., Denier van der Gon, J. J., and Gielen, C. C. (1982) Changes in recruitment order of motor units in the human biceps muscle. Exp Neurol 78, 360–368

